# Limited dispersal and an unexpected aggression pattern in a native supercolonial ant

**DOI:** 10.1101/2020.01.09.899971

**Authors:** S. M. Hakala, M. Ittonen, P. Seppä, H. Helanterä

## Abstract

Understanding how social groups function requires studies on how individuals move across the landscape and interact with each other. Ant supercolonies are extreme cooperative units that may consist of thousands of interconnected nests, and their individuals cooperate over large spatial scales. However, the inner structure of suggested supercolonial (or unicolonial) societies has rarely been extensively studied using both genetic and behavioral analyses. We describe a dense supercolony-like aggregation of more than 1 300 nests of the ant *Formica* (*Coptoformica*) *pressilabris*. We performed aggression bioassays and found that, while aggression levels were generally low, there was some aggression within the assumed supercolony. The occurrence of aggression increased with distance from the focal nest, in accordance with the genetically viscous population structure we observe by using 10 microsatellite markers. However, the aggressive interactions do not follow any clear pattern that would allow specifying colony borders within the area. The genetic data indicate limited gene flow within and away from the supercolony. Our results show that a *Formica* supercolony is not necessarily a single unit but can be a more fluid mosaic of aggressive and amicable interactions instead, highlighting the need to study internest interactions in detail when describing supercolonies.

## INTRODUCTION

Cooperation in social groups can be favored when interacting individuals are related, or otherwise share alleles for cooperative behavior (Hamilton 1964a, b). The balance between kin selected cooperation and harmful kin competition is a fundamental topic in the study of social evolution, and especially the roles of spatial patterns and dispersal have received considerable attention (e.g. Queller 1992; Taylor 1992; West et al. 2002; Kümmerli et al. 2009; Platt and Bever 2009). However, in natural populations, the spatial scales of cooperation and competition remain understudied. They can be especially difficult to interpret in genetically viscous populations, where limited dispersal leads to relatives aggregating together, leading to both increased kin competition and more possibilities for cooperation among relatives (Queller 1992; Taylor 1992). Understanding the relevant spatial scales of cooperation and competition requires knowledge on how individuals move across landscapes and interact with each other.

Ants are a good example of obligate eusociality, where a reproductive caste depends on help from a sterile worker caste (Crespi and Yanega 1995). In several ant species, colony structures have grown beyond the simple family unit, which is the ancestral state where eusociality originally evolved (Boomsma et al. 2014). In many ant taxa dispersal is limited, and new nests are formed by budding from a parent nest, instead of long range dispersal by flying sexual offspring (Cronin et al. 2013). In some of these taxa, this leads to the formation of polydomous colonies where several nests are interconnected and work together as a single colony (Debout et al. 2007). Further, polydomy is often connected to polygyny (multiple reproducing queens per colony) and the colonies can grow extremely large and have extremely high numbers of queens. This leads to colony members being unrelated at the local scale (Helanterä et al. 2009; Boomsma et al. 2014), which challenges traditional definitions of colonies as kin-selected cooperative units. Very large polydomous and polygynous colonies are often called supercolonies (Helanterä et al. 2009). In a supercolony, the cooperative colony spans larger spatial scales than a single individual can cross (Pedersen et al. 2006; Helanterä et al. 2009), but individuals still recognize and treat each other as members of the same colony when brought together (Moffett 2012). As an extreme example, a genetically homogenous single colony of the Argentine ant *Linepithema humile* has spread over the whole globe, but individuals living on different continents still behaved as one colony in behavioral experiments (Van Wilgenburg et al. 2010).

Although the existence of supercolonies is something of an evolutionary paradox due to low relatedness within these cooperative units (Giraud et al. 2002), they are ecologically very dominant. This has led to many supercolonial ant species becoming harmful invasive pests (GISD 2019), the most-studied example being the above-mentioned Argentine ant (Tsutsui and Case 2001; Giraud et al. 2002). It forms massive supercolonies especially in its invasive ranges, where it has been able to spread without much competition (Tsutsui and Suarez 2003; Wetterer et al. 2009), and smaller supercolonies in its native ranges (Pedersen et al. 2006; Vogel et al. 2009). However, supercoloniality has also evolved independently in several other taxa across the ant phylogeny (Helanterä et al. 2009). Studying the spatial scale and inner social organization of supercolonial societies in all of these taxa would give a fuller understanding of the evolution and maintenance of such high levels of cooperation (Robinson 2014).

The ability of ant individuals to distinguish between group members and outsiders makes it possible to define the borders of supercolonies (Moffett 2012). Individuals recognize their colony mates using olfactory cues that can be both genetically determined and acquired from the environment and food (Vander Meer and Morel 1998; Ginzel and Blomquist 2016). As ants usually behave aggressively toward intruders, aggression assays are commonly used to study nest and colonymate recognition, and provide a simple way to infer spatial scales of cooperation (Roulston et al. 2003). Even the largest argentine ant supercolonies lack inter-nest aggression within the colony, while they do behave aggressively toward other conspecific (super)colonies (Giraud et al. 2002; Moffett 2012). However, not all supercolonies and supercolonial species reported in the literature have been rigorously tested for inter-nest aggression or resource sharing (Hoffmann 2014), but instead some of them have been described as supercolonial based only on genetic data and the spatial organization of nests (Helanterä et al. 2009).

*Formica* ants offer excellent possibilities for studying spatial scales of cooperation and the evolution of social organization, as this genus has large variation in social organization, from simple family units all the way to very large supercolonies (Rosengren and Pamilo 1983; Rosengren et al. 1993; Helanterä et al. 2009; Ellis and Robinson 2014). The largest reported example is a 45 000 nest supercolony of *Formica yessensis* (Higashi and Yamauchi 1979), while supercolonies in other *Formica* species range from tens to a few thousands of nests (Markó et al. 2012). In some cases, populations of the same species vary in their social structure, and, for example in *Formica exsecta*, it seems that polygynous, polydomous, and supercolonial populations can arise from monogynous background populations (e. g. Seppä et al. 2004). *Formica* populations tend to be genetically viscous, i.e. spatially close nests are more closely related than spatially distant ones, which is especially true in polygynous species and populations where young queens are often philopatric (Rosengren et al. 1993; Chapuisat et al. 1997; Sundström et al. 2005). While patterns of genetic variation in supercolonial *Formica* ants have been studied in detail before (e.g. Chapuisat et al. 1997; Seppä et al. 2004, 2012; Elias et al. 2005; Sundström et al. 2005; Holzer et al. 2009), these studies have rarely combined genetic data with behavioral experiments to assess the scale of cooperation, or potential for it. Recognition behavior and inter-nest aggression has been tested in some species (Chapuisat et al. 2005; Holzer et al. 2006; Martin et al. 2009; Kidokoro-Kobayashi et al. 2012; Pohl et al. 2018), but overall the behavioral structure of highly polydomous *Formica* colonies remains understudied.

We investigate the nature and spatial scale of supercoloniality in the highly polydomous ant *Formica pressilabris* using behavioral assays and DNA microsatellite data. We test the hypothesis that behavioral colony borders correspond to the spatial and genetic structuring of a large nest aggregation, which is an underlying assumption of many previous studies. Based on this hypothesis, we expect nests at a densely populated *F. pressilabris* site to either belong to one supercolony without inter-nest aggression, or possible inter-nest aggression to occur between spatially or genetically distinct supercolonies competing with each other. We use genetic data to infer dispersal patterns within and outside of this supercolonial nest aggregation. As supercoloniality has previously been linked to limited dispersal, our hypothesis is that the dense nest aggregation is genetically somewhat separated from three other closely located study sites, and competition thus largely local. Additionally, we expect the philopatry of daughter queens to lead to genetic viscosity within our supercolonial site.

## MATERIAL AND METHODS

### STUDY SPECIES AND SITES

*Formica* (*Coptoformica*) *pressilabris* is a mound-building ant that lives on meadows and banks, builds nests of grass, and tends aphids for its main energy supply (Seifert 2000; Schultz and Seifert 2007). It founds new nests via temporary social parasitism with other *Formica* (*Serviformica*) species as its host, or via budding from a parent nest (Kutter 1969; Czechowski 1975). While monogynous colonies have been reported, secondary polygyny, where daughter queens stay in their natal nests, is common. A single nest can have hundreds of queens and grow up to over one meter in diameter, which is exceptionally large for such a small *Formica* species (Czechowski 1975; Collingwood 1979; Pamilo and Rosengren 1984; Rosengren et al. 1993; Seifert 2000). Colonies are also commonly polydomous with several interconnected nests and no aggression between nests (Czechowski 1975; Collingwood 1979; Seifert 2000). Nest turnover is high in polydomous colonies, i.e. new satellite nests are built regularly while old ones are abandoned (Bönsel 2007).

We sampled a *F. pressilabris* population in Southern Finland in 2016. The sampled area consists of one 9-ha abandoned field (Särkkilen) with a large continuous nest aggregation (hereafter referred to as the supercolonial site), and three closely located smaller fields with smaller nest aggregations (Storgård, Lillgård, Storsand) (Fig. 1). To assess whether the colonization potential of the area was fully used, we also extensively, although not exhaustively, searched for nests in other suitable habitats inside a one km radius from the supercolony site, but did not find additional colonies. Relatedness among worker nestmates estimated in a part of the supercolony site (0.21 +-0.02, Schultner et al 2014) indicates polygyny and/or polydomy, i.e. movement of individuals between nests. In our sampling, we counted nest mounds separated by more than 20 cm as separate nests. For our behavioral experiments and genetic analysis, we identified four spatially somewhat distinct parts of the supercolony site, with either more (parts II & III, Fig. 1), or less clear gaps between the parts (parts I & II, III & IV, Fig. 1). We calculated nest densities first for the whole supercolony site and then separately for each of the four parts on it as well as for the three smaller sites, using QGIS 3.4.1 (QGIS Development Team 2018). For the whole supercolony site, we used the complete open area of the field for the calculation, whereas for its four parts and the three separate fields we used the areas of the polygons obtained by drawing straight lines between the outermost nests belonging to the respective areas. The age of the study population is not known, but the supercolony site has not been cultivated since the 1970’s (landowner P. Forsbom, personal communication). Thus, the *F. pressilabris* population may have occupied the site for up to five decades.

**Fig. 1.**
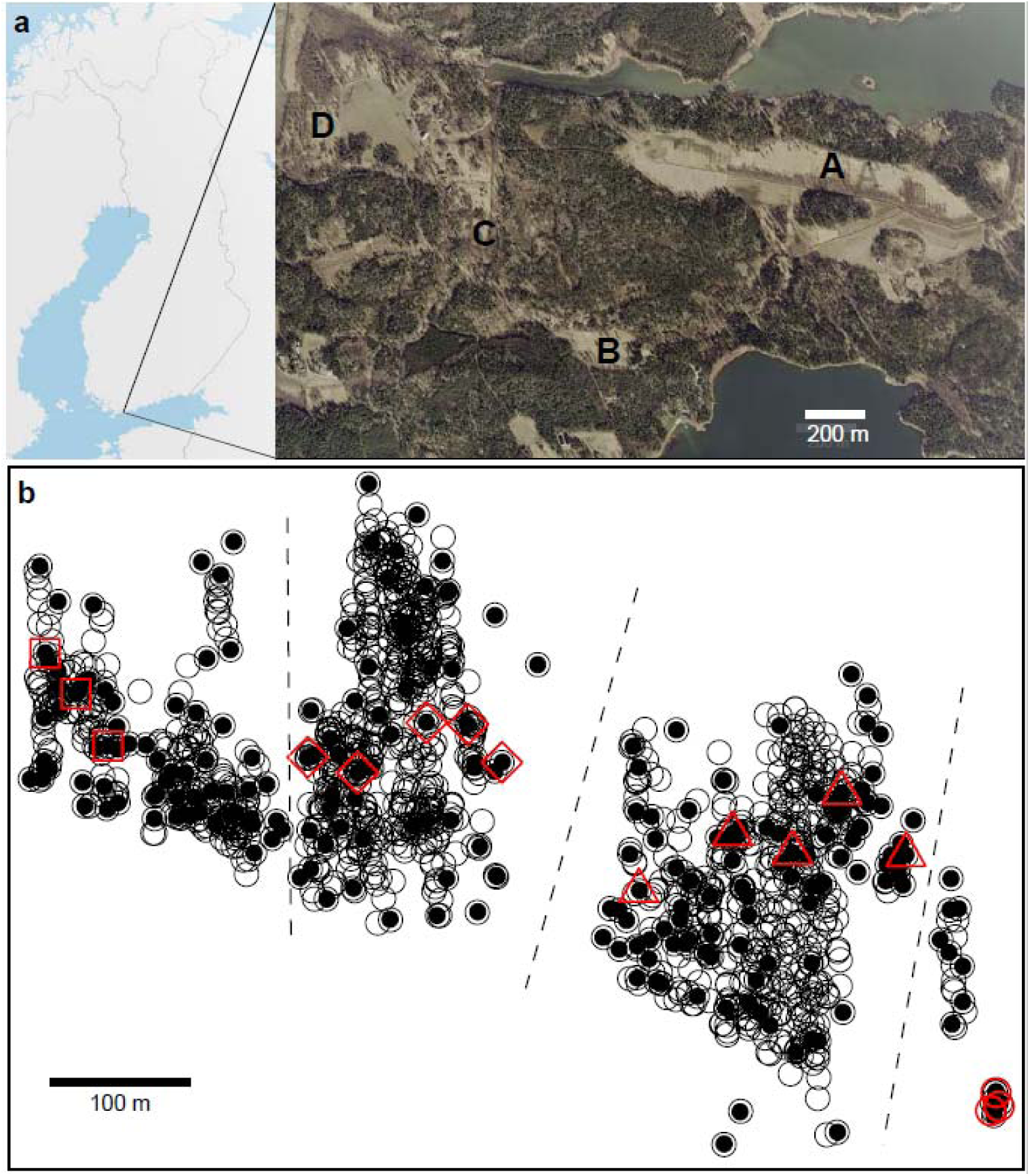
**a** The study area. The sampled subpopulations are indicated with letters. A: the supercolony field Särkkilen; B-D: the smaller fields, Storsand, Lillgård, and Storgård respectively. **b** Locations of the nests on the supercolony field (circles). The nests used for genotyping are marked with filled circles, and the nests used for the behavioral experiments with red symbols. The red squares, diamonds, triangles and circles represent parts I, II, III, and IV, respectively (see text). There are relatively large (> 30 m) nest-less gaps between these parts, except for parts I and II, between which there is a narrower area dominated by several *Formica exsecta* nests. Dashed lines mark the borders between the parts. The red circles in part IV have been slightly moved apart in order to show them all. Aerial image from National Land Survey of Finland NLS Orthophotos database 04/2019 (CC BY-SA 4.0).

### BEHAVIORAL ASSAYS

We performed behavioral assays in order to determine whether workers from the supercolony site behave differently toward their nest-mates, conspecifics from other close and distant nests at the same site, conspecifics from other sites, and allospecific ants. These treatments are hereafter referred to as control, neighbor, distant, outside, and allospecific, respectively. We expected no aggression within the supercolony site, and that if any aggression was observed, it would occur across the nestless gap in the middle of the site (Fig. 1), as the gap may form a barrier between two separate supercolonies. We expected more aggression toward individuals from other sites than toward individuals from within the supercolony site. The closely related species *F. exsecta* should always be faced with aggression, as *F. pressilabris* has previously been shown to behave very aggressively against it (Czechowski 1971).

We collected workers from 16 *F. pressilabris* nests at the supercolony site in June 2016, covering all four parts (Fig. 1). Additionally, we collected *F. pressilabris* workers from four nests at one of the smaller study sites, and one nest at another location approximately 30 km SW of the study population (Tvärminne). We also sampled two *F. exsecta* nests from the main study area and one from Tvärminne. We collected 2–5 liters of nest material and a minimum of 300 workers from each nest. We divided all 16 focal nests from the supercolony site into two boxes to control for the possibility that physical separation in the laboratory causes aggression in the assays. We reared the laboratory nests in room temperature, kept them moist and fed them daily with a Bhatkar-Whitcomb diet (Bhatkar and Whitcomb 1970).

We tested the reaction of workers from each of the 16 supercolony site nests against control, neighbor, distant, outside, and allospecific individuals (Fig 2.). We replicated each of the five treatments five times per nest with new arenas and new individuals, except for the allospecific tests. The latter we performed in the same arenas with the same individuals as the same-nest control treatments, as there never was any aggression in the control treatments. Our preliminary experiments showed that *F. pressilabris* workers act passively in standard one-on-one aggression assays on neutral arenas, usually showing no interest toward each other. Therefore, we used experimental arenas (diameter 6,5 cm) with 15 workers on their own nest material, simulating natural conditions with nest-mates and familiar odors present. In an assay like this, the observed behavior is expected to correlate with the natural nest defense behavior, revealing whether the workers would allow visitors to enter the nest or not, as even submissive ant species defend their nests against intruders (Vepsäläinen and Pisarski 1982; Savolainen and Vepsäläinen 1988). If an assay this sensitive does not show aggression, this can be interpreted as very strong potential for cooperation: when ants are willing to share their nest, they likely cooperate in other ways, too. After letting the 15 focal ants calm down (when they had stopped running around and did not show signs of alert, such as opening their mandibles), we introduced one worker from another laboratory nest box and recorded the actions of the ants for one minute, using a Canon EOS 550D DSLR camera with a Canon EF 100mm f/2,8 macro lens. The distance between the ants and the lens was 48.5 cm. Ants did not react toward the camera. We performed the behavioral assays on 3–7 July 2016.

**Fig. 2.**
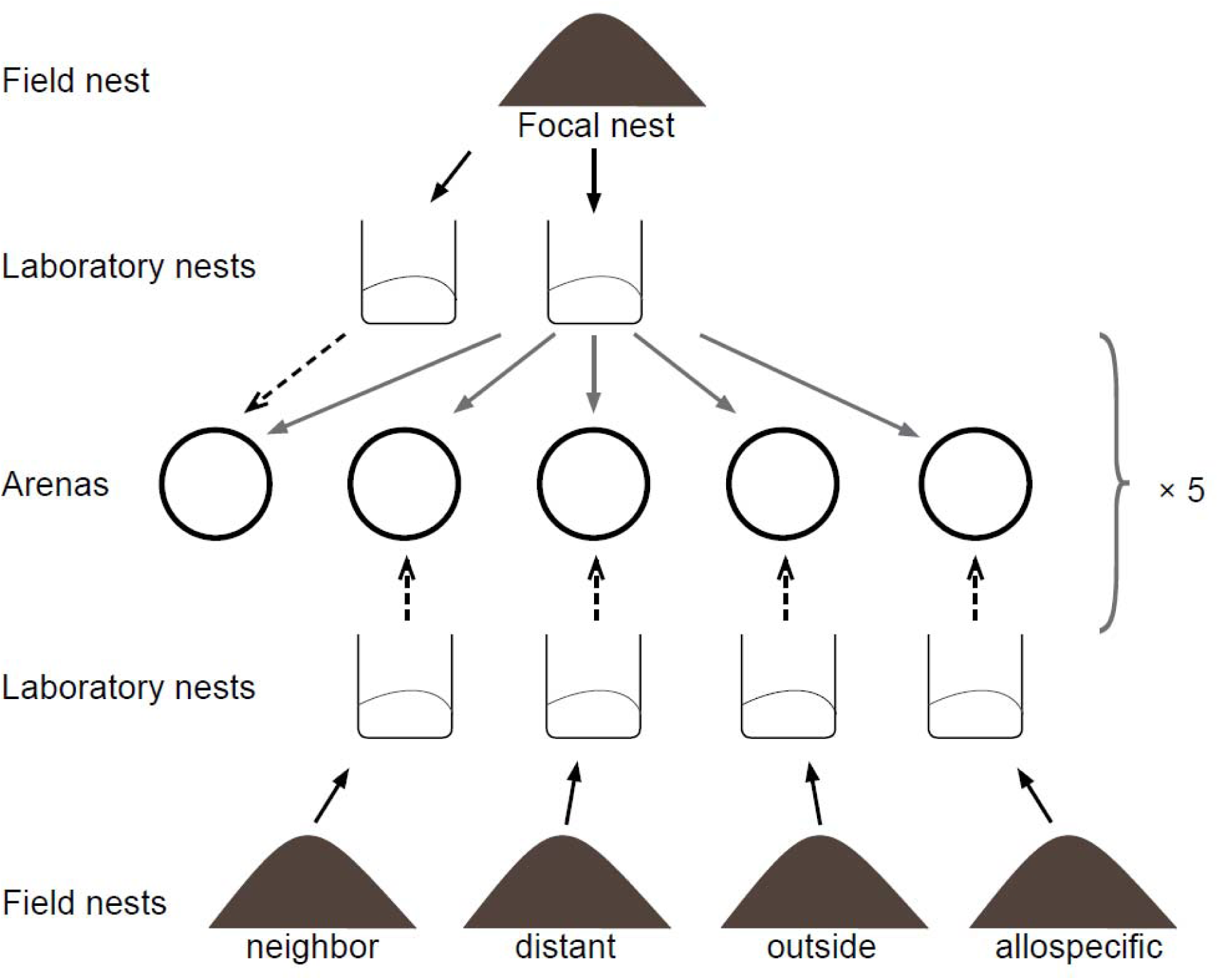
The design of the behavioral experiment. In each assay, we tested the reactions of 15 ants from a focal nest toward one ant from another laboratory nest box. We collected ants and nest material into laboratory nest boxes (black arrows), and put 15 workers from a focal nest into each of five experimental arenas (grey arrows). Then we introduced one worker from another laboratory nest into the arenas (dashed arrows). For each focal nest, we replicated this procedure five times with new ants and arenas. Control = introduced ant from the same nest, neighbor = introduced ant from the same part of the supercolony field, Distant = introduced ant from a different part from the supercolony field, Outside = introduced ant from another field, Allospecific = introduced ant of a different species, *F. exsecta*

One of the authors (MI) watched and transcribed the videos in a randomized order at half speed and blindly regarding which nests and which treatments were represented in each video. The durations of antennation, trophallaxis and biting events against the one introduced worker were recorded using the software JWatcher 1.0. Mandible opening and other minor signs of aggression could not be reliably scored due to the small size and large number of individuals on the arena. The scoring was hierarchical in the sense that when even one of the fifteen focal individuals was aggressive, we did not score nonaggressive behavior at the same time, because aggression-related pheromones could affect the behavior of other individuals. In our analyses, we combined trophallaxis and antennation as nonaggressive inspecting behavior, because the two were sometimes hard to separate from each other and both have been shown to increase when individuals recognize the opponent as a non-nestmate in polydomous *Formica paralugubris* (Chapuisat et al. 2005; Holzer et al. 2006). Seven out of the 400 videos could not be analyzed due to damaged files.

### STATISTICAL ANALYSIS OF BEHAVIORAL DATA

We analyzed the presence and absence of aggression explained by the different treatment classes with a binomial generalized linear mixed model (GLMM). In all of our models, we included both the nest of the focal workers and the nest of the introduced worker as random effects to account for the nonindependence of samples coming from the same nest. We excluded the ‘control’ treatment as no aggression occurred in the within-nest controls. We used a beta GLMM to analyze the duration of aggression among the treatment classes where it occurred. In the videos where no aggression occurred, we also used a beta GLMM to analyze the duration of nonaggressive inspecting behavior (antennation and trophallaxis combined). For this analysis we excluded the treatment level ‘allospecific’ due to low sample size (n = 3), and substituted five samples with a value of 0 (= no inspecting behavior) with a value of 1 (= a millisecond of inspecting behavior) to allow the use of the beta distribution, which cannot contain zeros. For the beta GLMM’s, we measured the duration of aggression or inspecting behavior as the proportion of the total time that the introduced ant was in sight.

We further tested whether the geographical distance between two separate nests explains the presence or duration of aggression, or the duration of nonaggressive inspecting behaviors. We analyzed a subset of our behavioral data within the supercolony site (treatment levels ‘neighbor’ and ‘distant’), using a binomial GLMM for presence of aggression and beta GLMM for duration of aggression and inspecting behavior. Geographical and genetic distances are collinear in our data (see below) and the effect of these two variables cannot be fully separated in our results. Therefore, we used only geographical distance as an explanatory variable in our analysis. As we did the genetic analysis only for a single individual per nest, using genetic distance would be more problematic in connection to the nest-level behavioral data. Additionally, conspecific aggression in ants correlates mostly with chemical distance (Martin et al. 2012) that may have both genetic and environmental components (Vander Meer and Morel 1998; Ginzel and Blomquist 2016). Thus, as geographic distance contains information of both genetic and environmental factors, we deemed it more biologically relevant than genetic distance. We analyzed the behavioral data in R (R Core Team 2013) with the package glmmTMB (Bolker et al. 2009).

### MICROSATELLITE GENOTYPING AND POPULATION GENETICS

To estimate gene flow between and within the four sites, we genotyped a single worker from 285 different nests, including 213 nests from the supercolony site (Fig. 1) and all nests from the three smaller study sites. We extracted the DNA using NucleoSpin Tissue extraction kits (Macherey-Nagel) and genotyped the samples with 14 microsatellite markers originally developed for other Formica species (Chapuisat 1996; Gyllenstrand et al. 2002; Trontti et al. 2003; Hasegawa and Imai 2004) using the protocol designed by Hakala et al. (2018). We scored the microsatellite alleles with the software GeneMapper 5.

We analysed linkage disequilibrium and Hardy-Weinberg equilibrium using GenePop on the Web (Raymond and Rousset 1995; Rousset 2008) and calculated allelic richness values using the PopGenReport 3.0.0 package (Adamack and Gruber 2014) in R. We calculated allele frequencies, linear genetic distances between nests, and heterozygosity values using GenAlEx 6.502 (Peakall and Smouse 2012). We calculated F_ST_ values among all sampled sites, and among the four parts within the supercolony field using AMOVA’s in GenAlEx 6.502 (Peakall and Smouse 2012). To test for genetic viscosity, we performed Mantel tests using the package ecodist 2.0.1 (Goslee and Urban 2007) in R.

We analyzed the genetic structure of the population with a Bayesian approach using the software STRUCTURE 2.3.4 (Pritchard et al. 2000), which clusters individual genotypes by probability of similarity. To determine the most likely number of clusters (K) the analysis was run with K ranging from 1 to 7, using the admixture model and correlated allele frequencies. For each K value, we ran the analysis ten times with a burn-in of 100,000 steps for a run length of 300,000 steps. We estimated the most likely number of clusters by applying the delta K method with plotting the mean and standard deviation of the mean likelihood L(K) for each run in STRUCTURE HARVESTER (Evanno et al. 2005; Earl and vonHoldt 2012). As the mathematical model used by STRUCTURE is not ideal with unbalanced sampling and groups with low sample sizes or patterns of isolation by distance (Kalinowski 2011; Puechmaille 2016), we repeated the genetic mixture analysis with a similar software, BAPS 6.0 (Corander et al. 2003; Corander and Marttinen 2006; Corander et al. 2008). BAPS was allowed to find the most probable number of clusters with repeated runs (10 times K1-K7). As BAPS was unable to find any stable clustering without a spatial prior, we repeated the analysis as a spatial analysis with geographical coordinates. Subsequently, we performed an admixture analysis with the results of the spatial analysis.

## RESULTS

### MAPPING OF THE STUDY AREA

The supercolony site had more than 1,300 nests (Fig. 1, Appendix table A1), and the nest densities ranged from 254 to 401 nests/ha in the different parts of the site. The three smaller sites had 7, 16, and 27 nests, and the nest densities were 426, 25, and 23 nests/ha, respectively. We could not directly observe worker movement between nests, because *F. pressilabris* workers walk mostly on the ground surface under the grass cover or on grass stems, forming no visible paths between the nests. All parts of the supercolony site had many dense aggregations with nests situated very closely together. Often these aggregations had one or two large main nests and a few smaller ones. The distances between nest aggregations were often short, and the field is overall almost uniformly occupied by the species.

### BEHAVIORAL ASSAYS

There was no aggression in the same nest controls, while the allospecific treatment with *F. exsecta* had aggression in 72 out of 75 assays (Fig. 3). In the neighbor, distant, and outside treatments there was clearly more aggression than in the completely nonaggressive control treatments. Aggression occurred significantly more often in the allospecific than in any other treatment (compared to Neighbour: Z=6.71, SE=0.73, p<0.001; Distant: Z=5.64, SE=0.71, p<0.001; Outside: Z=5.08, SE=0.72, p<0.001). The workers were also more often aggressive toward conspecific ants from distant nests than those from neighbor nests (Z=2.23, SE=0.40, p=0.026). However, there was no significant difference between the aggression faced by ants from distant and outside nests (full test statistics in Appendix table 2). The behavioral patterns are not consistent among the five replicates, but instead there are plenty of nonaggressive replicates also between distant nest pairs. (Fig. 4).

**Fig. 3.**
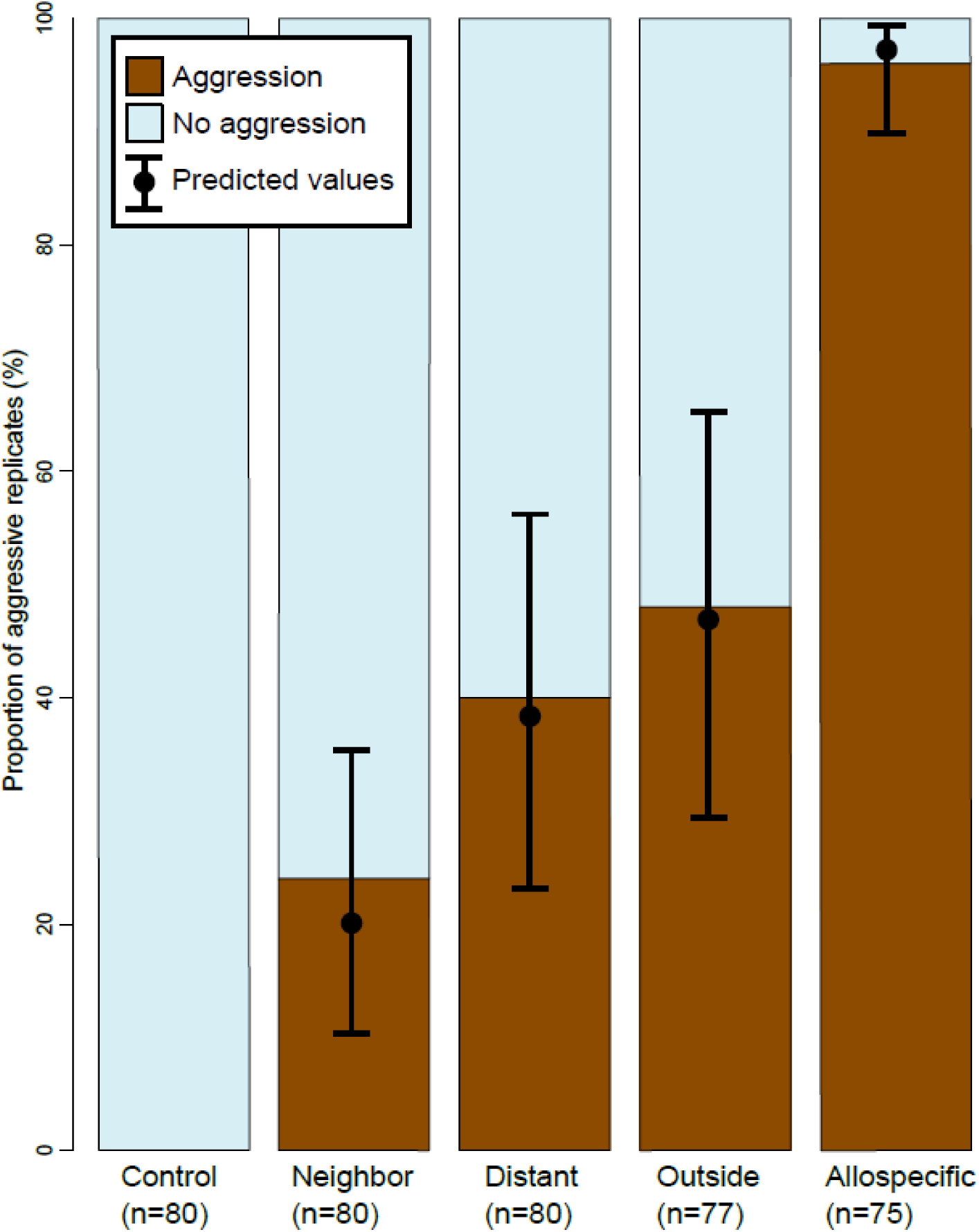
Presence and absence of aggression by treatment with model predictions for the four treatment classes that were included in the analysis (binomial GLMM). All other differences were significant, except the difference between distant and outside. Control = introduced ant from the same nest, neighbor = introduced ant from the same part of the supercolony field, Distant = introduced ant from a different part from the supercolony field, Outside = introduced ant from another field, Allospecific = introduced ant of a different species, *F. exsecta*

**Fig. 4.**
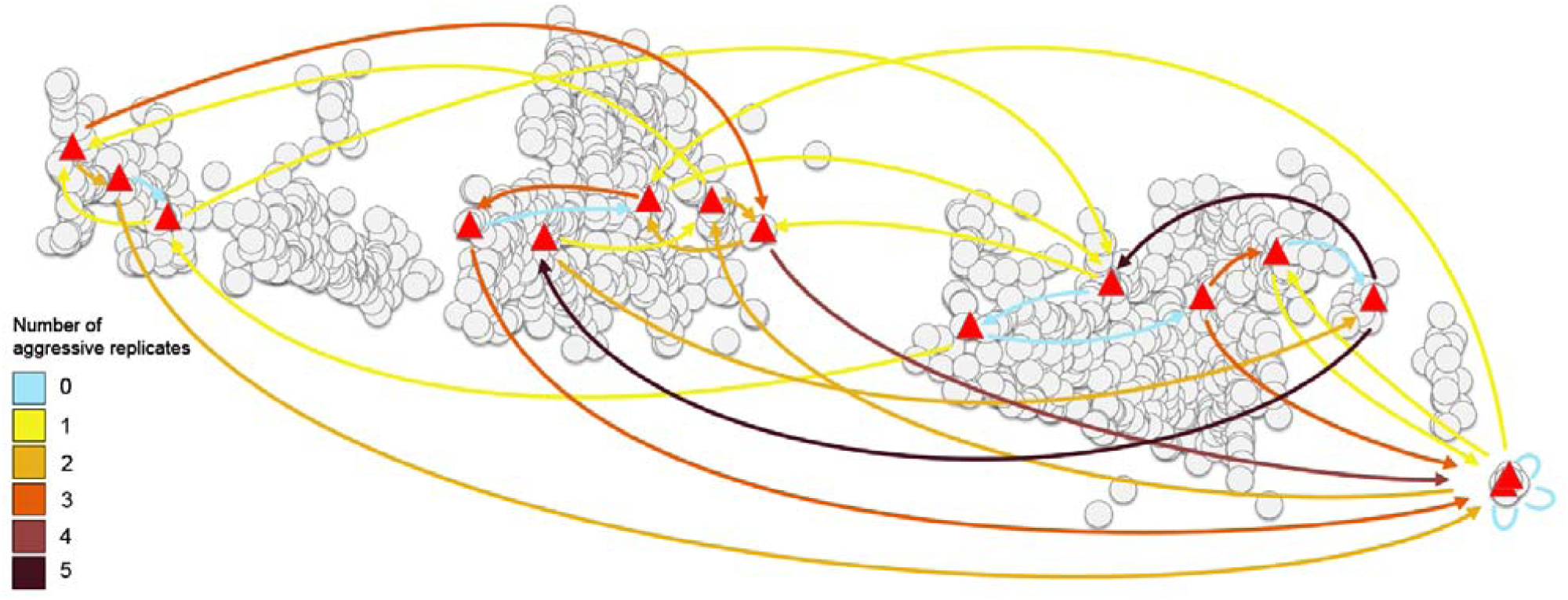
The number of replicates (out of five) with aggression for each nest pair within the supercolony site. The nests used in the aggression assays are shown as red triangles, other nests as grey circles. The colored arrows show the number of aggressive replicates (see legend), and arrowheads show the direction of aggression, pointing toward the nest of the introduced ant.

In the assays where aggression occurred, its duration did not significantly differ among the treatments, except in one of the pairwise comparisons, where the effect size remained small (Fig. 5). However, the allospecific treatment always had long aggression durations (>25% of the assay time), whereas all of the within-species treatments also had some short durations (<25% of the assay time). In the assays without aggression, significantly less inspecting behavior was targeted toward the control than any of the other treatments (compared to Neighbour: Z=5.45, SE=0.19, p<0.001; Distant: Z=5.07, SE=0.20, p<0.001; Outside: Z=3.74, SE=0.34, p<0.001), while none of the other treatments differed from each other (Fig. 6, Appendix table 2).

**Fig. 5.**
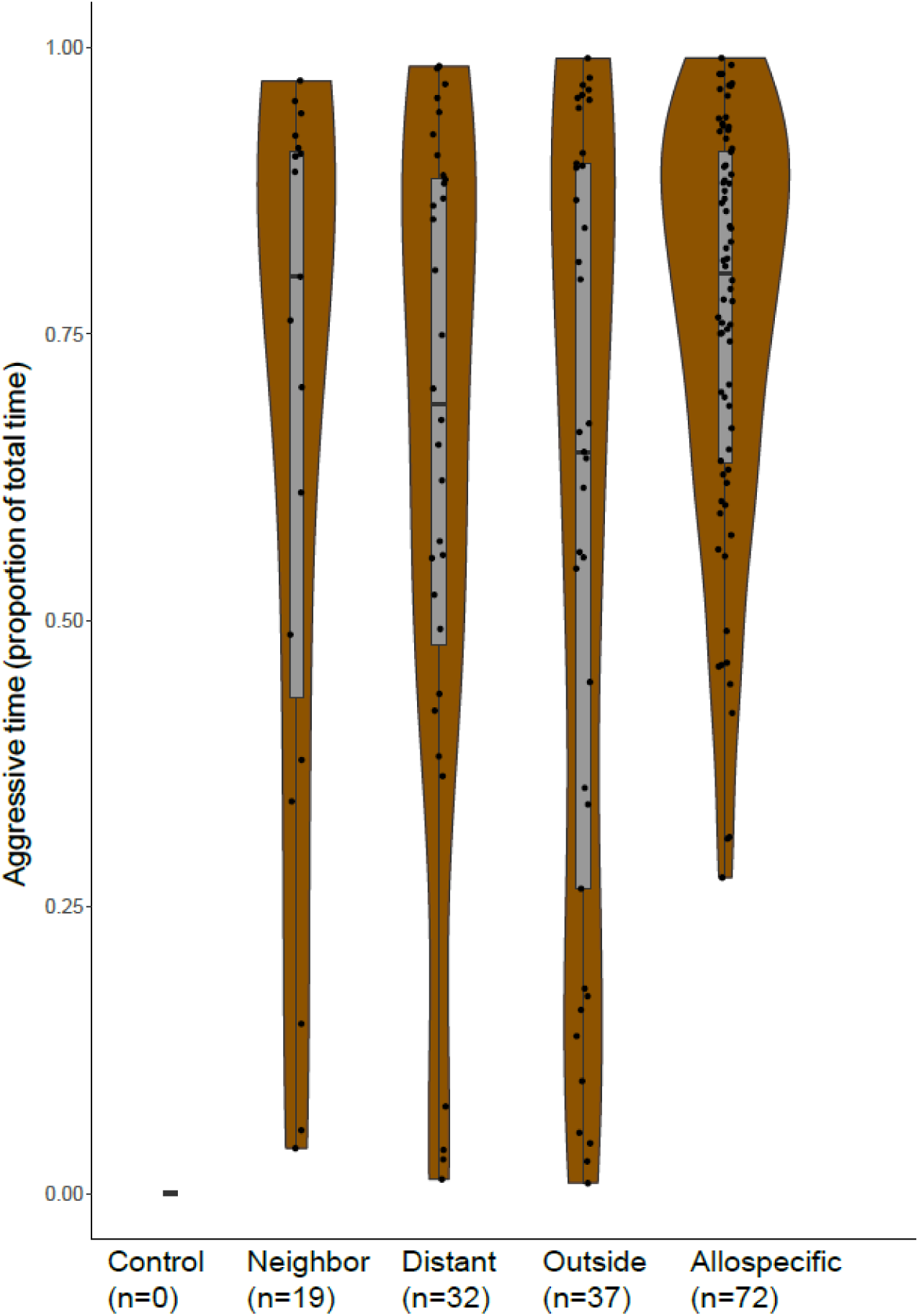
Duration of aggression (when present) by treatment as proportion of the total observation time. All data points, density plot, and median and quartile plot (box plot) shown. Only the difference between ‘outside’ and ‘allospecific’ is statistically significant (beta GLMM: X=2.43, SE=0.23, p=0.015). Control = introduced ant from the same nest, neighbor = introduced ant from the same part of the supercolony field, Distant = introduced ant from a different part from the supercolony field, Outside = introduced ant from another field, Allospecific = introduced ant of a different species, *F. exsecta*.

**Fig. 6.**
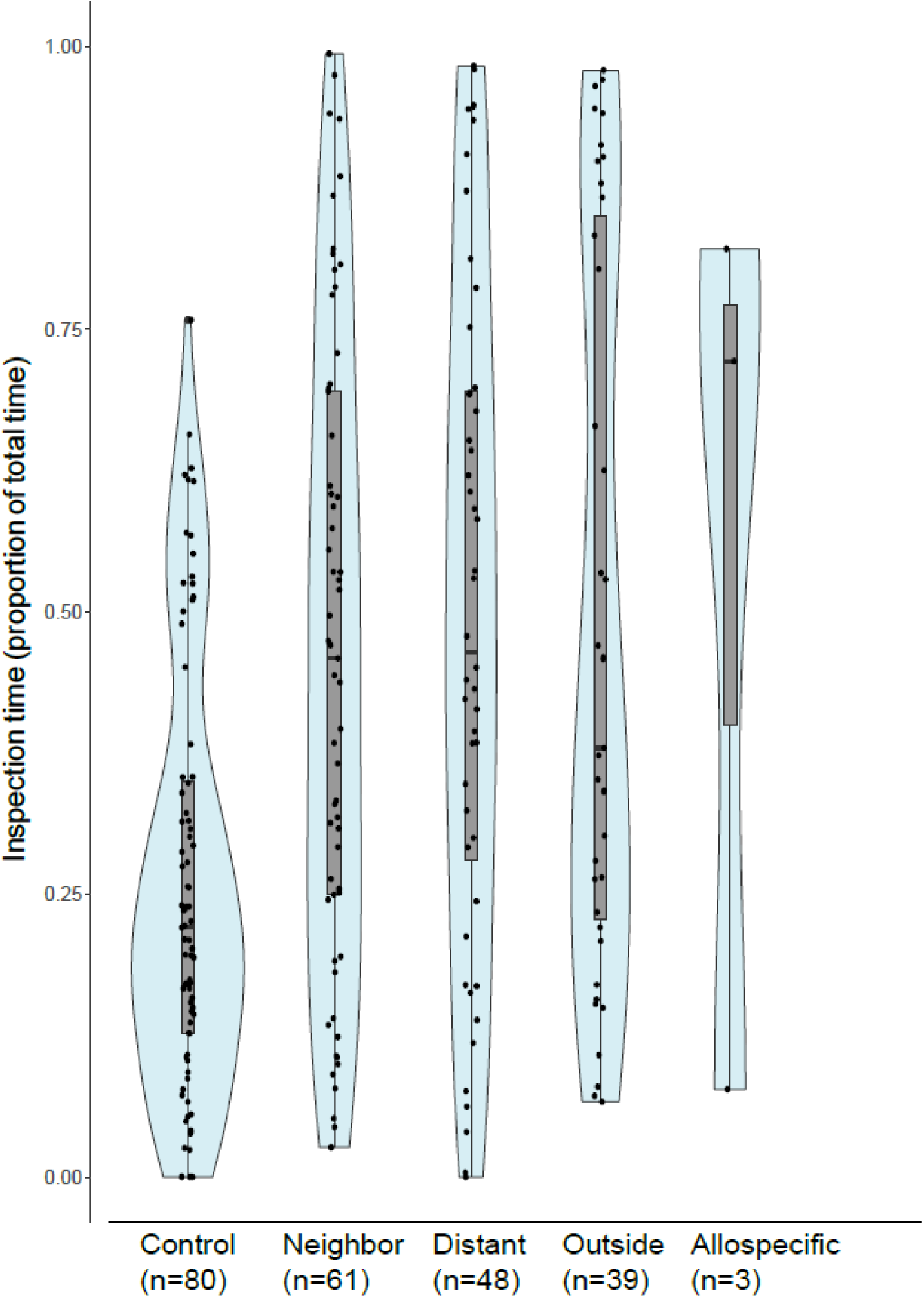
Duration of inspection behavior (antennation and trophallaxis) by treatment in the nonaggressive samples as proportion of the total observation time. All data points, density plot, and median and quartile plot (box plot) shown. Only the treatment Control is significantly different from other classes (beta GLMM, allospecific treatment not included in the model due to n = 3). Control = introduced ant from the same nest, neighbor = introduced ant from the same part of the supercolony field, Distant = introduced ant from a different part from the supercolony field, Outside = introduced ant from another field, Allospecific = introduced ant of a different species, *F. exsecta*.

Within the supercolony site, the occurrence of aggression between two nests increased with geographical distance (Fig. 7a, binomial GLMM, z = 2.85, SE=1.20, P = 0.004), but its duration did not change with distance (Fig. 7b, beta GLMM z= -0.334, SE=0.94, P = 0.74). The duration of inspecting behavior between nests within the supercolony field did not change with increasing distance (beta GLMM, z= 0.36, SE=0.66, P = 0.72).

**Fig. 7.**
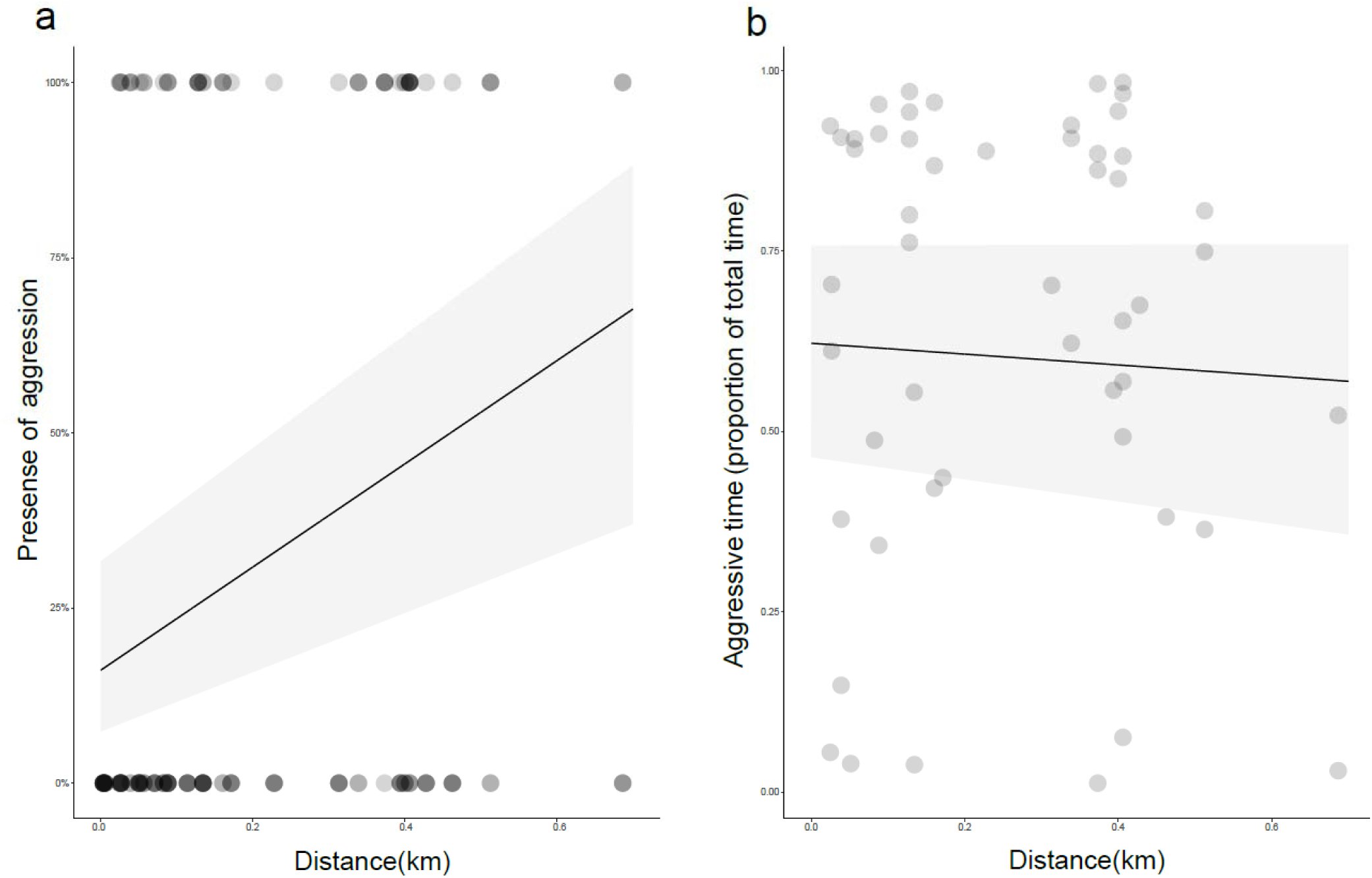
**a** Presence of aggression between different nests on the supercolony field as a function of internest distance, data (dots) and prediction (line with 95% confidence intervals) according to a binomial GLMM). N= 160. Darker dots indicate several data points on top of each other. **b** Duration of aggression (when present) between different nests as a function of distance on the supercolony field, data (dots) and nonsignificant prediction (line with confidence intervals according to beta GLMM). N=51

### POPULATION GENETICS

Of the original 14 microsatellite markers, we used 10 for further analysis, as four had too much missing data or significant heterozygote deficiency at all study sites (details in Supplementary material). The pairwise F_ST_ values show that the four parts of the supercolony site differed genetically from each other and that this differentiation was on a level similar to the differentiation among the other study sites (Table 1). Mantel tests showed minor but significant genetic viscosity when analyzing all samples in the study area (R = 0.06, 95% CI = 0.04, 0.08; P = 0.038), and also when analyzing only the samples from the supercolony site (R = 0.1, 95% CI = 0.09, 0.12; P ≤ 0.001).

**Table 1.**
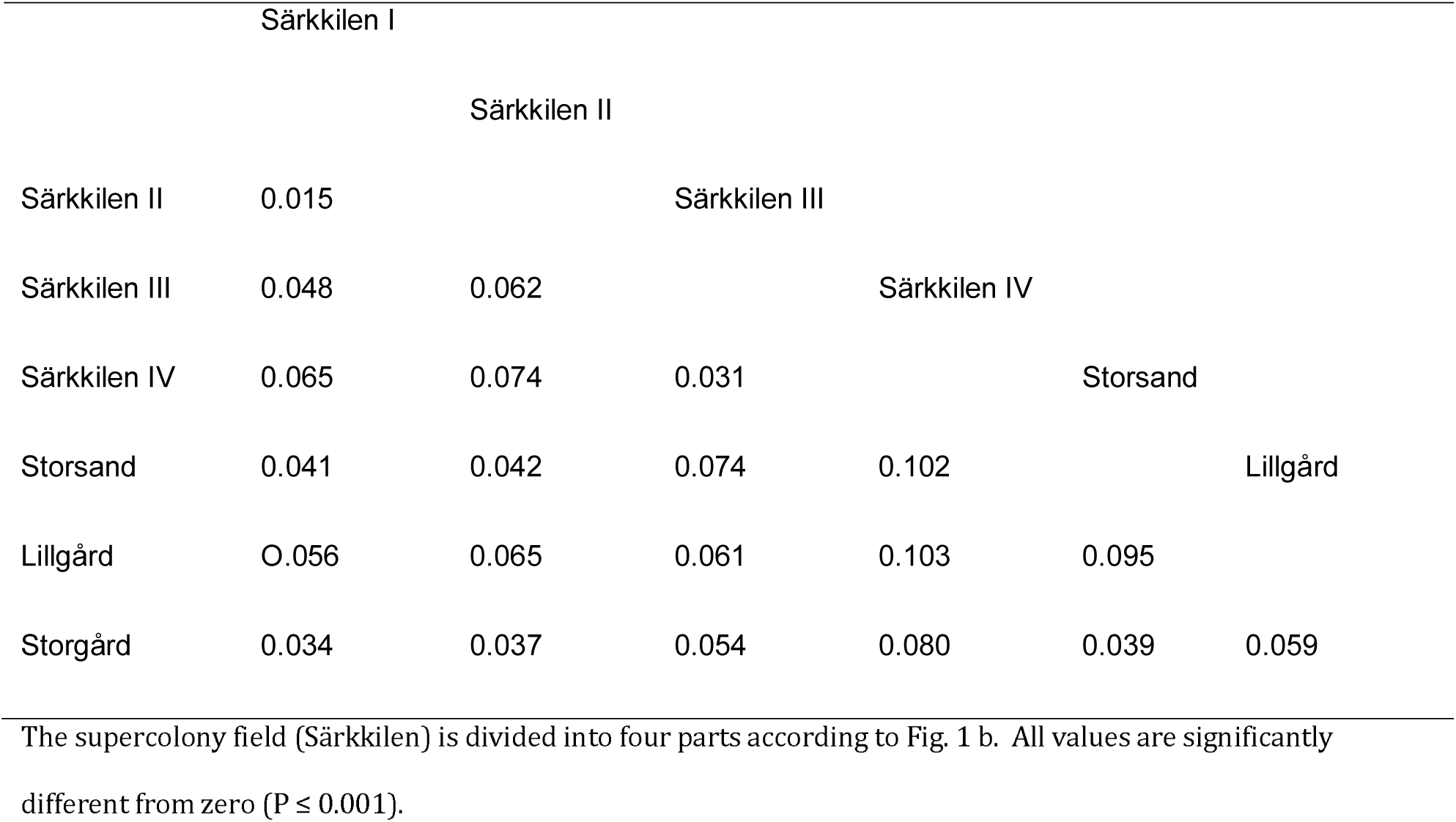
Pairwise F_ST_ values between subpopulations from an AMOVA with 999 permutations.

In the Bayesian clustering for the study area (Fig. 8 a&b), the optimal number of genetic clusters was two according to STRUCTURE, and three according to BAPS (posterior probability = 0.981). However, the obtained clusters did not correspond to the different locations, as both analyses showed some sites to contain a mixture of individuals belonging to different clusters. The results from STRUCTURE revealed strong genetic admixture among individuals, whereas BAPS found admixture only in a few individuals. The four sampled sites seem to belong to a single population with gene flow among the sites.

**Fig. 8.**
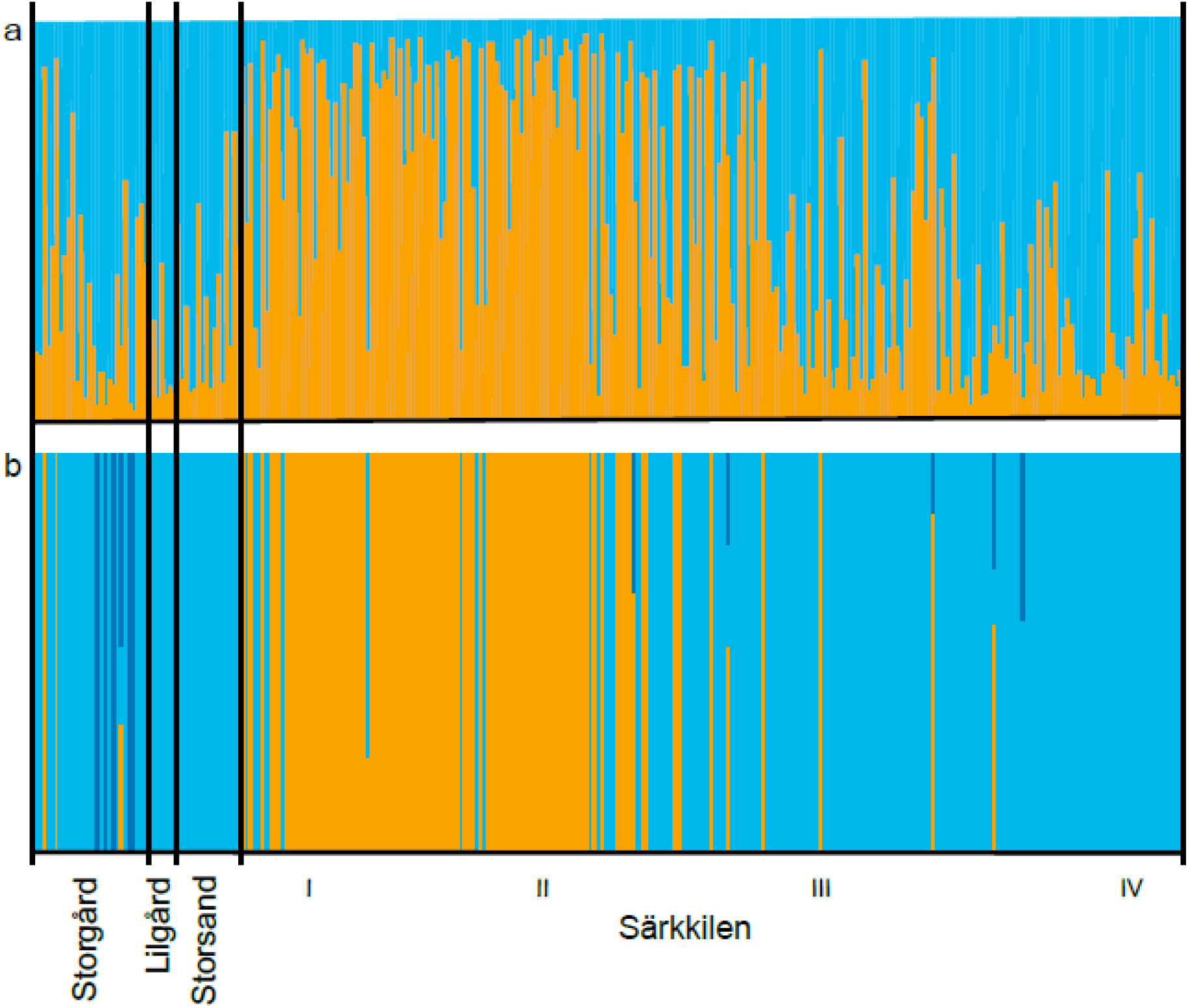
**a** Bayesian clustering of all genotyped individuals from the four study sites (I-IV = the four different parts of the supercolony site) with the software STRUCTURE. Samples organized from west to east. Optimal K = 2 with strong admixture **b** Bayesian clustering of the same data with the software BAPS. Optimal K = 3 with less admixture

## DISCUSSION

We found minor genetic viscosity on a small spatial scale, both within the supercolonial site and in its close surroundings. This indicates limited dispersal within the study area, as expected if new nests are formed mostly by budding from parent nests. However, the fact that the different sites are not genetically more differentiated than different parts of the supercolony site indicates that some dispersal by flight also occurs. Behavioral experiments within the densely populated supercolony site show a similar behavioral pattern: the overall aggression level is low, workers mostly tolerate visitors from other nests in conditions simulating their own nest environment, and geographically close nests are better tolerated than nests further away. This suggests potential for cooperation among adjacent nests, and this potential slightly decreases the further nests are apart. However, because we observed some aggression within the supercolony site, it does not seem to consist of a single, distinct supercolony. There might be a mosaic of multiple supercolonies at the site, but as our data do not reveal any clear-cut behavioral borders, it is possible that the different colonies are somewhat connected over the whole spatial scale.

### GENE FLOW AND DISPERSAL

Our population genetic data indicate limited dispersal within the supercolony site. Even though within-nest relatedness is low, probably due to polygyny and mixing of individuals among adjacent nests (Schultner et al. 2014), individuals do not seem to mix effectively across the entire supercolony site, not even winged sexual individuals. Such genetic viscosity at a site less than one kilometer across shows that these ants disperse mostly over very short distances. Local mating between nestmates or individuals from neighboring nests must be common, as the population would otherwise not remain even weakly genetically viscous. *F. exsecta*, which is closely related and ecologically similar to *F. pressilabris*, has similar levels of genetic viscosity in its supercolonies (Seppä et al. 2012). In *Formica paralugubris*, a supercolony was more viscous than the surrounding non-supercolonial population, which, just as our results, suggests that supercoloniality is linked to locally reduced dispersal (Chapuisat et al. 1997). Importantly, data on gene flow, such as in the above-mentioned cases, do not provide any information about failed dispersal attempts. Workers in existing nests have a key role in determining which sexual individuals can establish themselves as reproducers at the supercolony site. Aggressive behavior toward individuals from distant nests could make their establishment as reproducers hard, even if they tried.

On a slightly larger spatial scale, the supercolony site is not genetically distinct from surrounding smaller polydomous colonies. Instead, pairwise F_ST_ values among the small sites and among the different parts of the supercolony site are in the same range. Our data suggest that longer-range dispersal among different sites is frequent enough to keep population structuring low, although dispersal seems to be limited within the supercolony site. Ongoing long-range dispersal ensures the supercolony is not a closed a population, and extends the scale of competition beyond single sites, which may give some selective advantage for the individuals from the supercolony on larger spatial scales (Pedersen et al. 2006; Kennedy et al. 2014). This contrasts with previous findings in F. exsecta, where polydomous colonies were genetically more different from surrounding monodomous and monogynous colonies than these were from each other (Seppä et al. 2004; Gyllenstrand et al. 2005), suggesting that polydomous colonies could form closed populations without much dispersal outwards.

Unfortunately, with our data we cannot assess whether dispersal is sex biased. Male biased dispersal is common in polygynous ants (Hakala et al. 2019) and in *Formica* overall (Sundström et al. 2005). In socially parasitic *Formica* species, such as our study species, queen dispersal is further complicated by the fact that queens cannot found their nests alone, but have to parasitize a nest of their host species or possibly a foreign nest of their own species (Czechowski 1975; Buschinger 2009). Reduced queen dispersal, or queen dispersal predominantly among existing colonies instead of founding new ones, would impair the colonization potential even if dispersal abilities were good, as suggested by gene flow among sites. Indeed, there are plenty of empty potential habitat patches around the supercolony in our study area, which speaks for limited colonization. Overall, the dispersal strategy of *F. pressilabris* seems two-fold: risky long-range dispersal combined with high levels of queen philopatry and potential to spread locally through short-range dispersal by foot. Such a dual strategy seems to be the rule in polygynous and polydomous *Formica* (Sundström et al. 2005) and exists in other ant taxa too, e.g. in *Crematogaster pygmaea* (Hamidi et al. 2017).

### AGGRESSION PATTERNS AND POTENTIAL FOR COOPERATION

Our main study site is, to our knowledge, the largest described nest aggregation of *Formica pressilabris* and among the largest of any species in the *Coptoformica* subgenus (Czechowski 1971, 1975; Markó et al. 2012). The nest densities (Appendix table A1) are well in line with previously reported values for large polydomous systems of *Formica* (Markó et al. 2012). While we are not aware of any reported nest densities for monodomous *F. pressilabris*, Pamilo and Rosengren (1984) reported clearly lower values (1.16–3.11 nests/ha) for three monodomous *F. exsecta* populations. The nest densities and observed high tolerance for introduced workers make us confident that the supercolony field is polydomous to a large degree. However, our behavioral assays still suggest that this nest aggregation is not a uniformly cooperative supercolony. Aggression generally increases with distance, but there are plenty of exceptions and no distinguishable colony borders (Fig. 4). This behavioral pattern resembles a social equivalent of a ring species, where all individuals that meet each other in natural settings may interact peacefully, and all of the nests can thus be considered to belong to one colony, but individuals may act aggressively when experimentally brought together from distant parts of the range (Moffett 2010; Moffett 2012).

In contrast to our data, dense and spatially distinct polydomous nest aggregations often show a complete lack of aggression, e.g. in both native and introduced argentine ants (Giraud et al. 2002; Björkman-Chiswell et al. 2008; Vogel et al. 2009), introduced *Myrmica rubra* (Chen et al. 2018), and native *Formica* (Chapuisat et al. 2005; Holzer et al. 2006; Kidokoro-Kobayashi et al. 2012; Pohl et al. 2018). However, some previous studies have also shown aggression within large polydomous *Formica exsecta* colonies (Pisarski 1982; Katzerke et al. 2006). Observations of seasonal and resource dependent differences in aggression levels (Mabelis 1979; Mabelis 1984; Sorvari and Hakkarainen 2004), and seasonal variation in supercolony genetic structure (Elias et al. 2005; Schultner et al. 2016), suggest that further temporal analysis of aggression patterns in supercolonial *Formica* is needed.

In addition to the current study, positive correlations between aggression and spatial distance have been reported e.g. within polydomous sites of *F. exsecta* (Katzerke et al. 2006) and *Myrmica rubra* (Garnas et al. 2007; Fürst et al. 2012), although later Chen et al. (2018) did not find such a correlation in *M. rubra*. Importantly, Chen et al. (2018) had, through a previous set of aggression assays, assigned colony borders prior to testing for this relationship, and suggest that the correlations between geographic distance and aggression found by Garnas et al (2007) and Fürst et al. (2012) may be attributed to mixing nest pairs belonging to the same and different colonies in one analysis. Our supercolonial site may indeed consist of multiple supercolonies instead of one, but the lack of clear patterns and the overall low levels of aggression revealed in our study (Fig. 4) lend little support to the existence of clear and persistent borders at our study site. Finally, aggression has been shown to increase with inter-nest distance also in monodomous species (Beye et al. 1998) and among distinct colonies or sites (Rosengren et al. 1986; Pirk et al. 2001; Holzer et al. 2006; Zinck et al. 2008), making these kinds of behavioral patterns hard to interpret.

The different behavioral assay methods used in many of the studies discussed above make direct comparisons of the results difficult (Roulston et al. 2003). Our method, where 15 individuals on their own nest material met one introduced ant, is very sensitive to aggression as it simulates an alien ant suddenly appearing in a nest, and there are many ants that can react. Based on our pilot experiments we consider it likely that *F. pressilabris* would have shown even less, if any, aggression in some more commonly used assay types, such as one-on-one tests on neutral arenas. Even in the absence of aggression, our results show that workers spend more time inspecting any nonnestmates than nestmates. Thus workers may distinguish between nestmates and more distant individuals, as is also suggested by increased antennation and trophallaxis in *F. paralugubris* (Chapuisat et al. 2005; Holzer et al. 2006), and increased antennation in argentine ants (Björkman-Chiswell et al. 2008).

While a lack of aggression between nests is commonly interpreted as a sign of shared colony identity, it does not necessarily mean that two nests share resources (Giraud et al. 2002; Heller et al. 2008; Buczkowski 2012). In large nest aggregations it is relevant to ask what the true spatial scale of cooperation and competition is (Pedersen 2012). At our supercolony site, aggression did not always occur even when testing over the nestless gap in the middle of the field, showing that workers are willing to let individuals from very far away enter their nests (Fig. 4). However, we consider true cooperation over this gap unlikely: given the width of the gap (∼70 m), worker movements over it should be rare. Czechowski (1975) found that workers can move at least 20-30 meters between nests in supercolonies of *F. pressilabris*, but such movements were considerably rarer than movements between nearer nests. In *F. exsecta*, workers from polydomous colonies forage on trees less than ten meters from a central nest (Sorvari 2009). If resources are shared over relatively limited distances also at our study site, workers of the two field halves belong to functionally separate colonies even though they are not clearly distinct based on either genetic or behavioral data. We agree with Lester and Gruber (2012) in their argument that functional cooperation and resource sharing is a crucial component when considering the evolution and maintenance of supercolonies. To be able to assess true relatedness among cooperating individuals we need to understand which parts of assumed supercolonies truly cooperate, and whether there are seasonal or resource dependent patterns. Without this knowledge, it is not possible to assess whether competition happens more within or among assumed supercolonies.

The genetic viscosity corresponds to the behavioral pattern where workers from nearby nests were allowed to enter the nest material more than distant workers. This suggests that limited dispersal does result in cooperation among relatives in *Formica* supercolonies. As our genetic data suggests that competition over reproduction is not exclusively local, local cooperation even under low but positive relatedness may help maximizing reproductive success on a larger spatial scale. The *F. pressilabris* nest aggregation described in this study is extremely dense and seemingly supercolonial. However, it defies usual definitions of ant colonies as single cooperative units with clear borders. Based on our behavioral data, discrimination in *F. pressilabris* is fluid, which begs for further studies on the functional connectedness of the nests, and the cooperative behavior in more natural settings in the field. Truly understanding the nature of supercoloniality requires more functional studies focusing on resource sharing and competitive dynamics—in all ant taxa exhibiting this fascinating lifestyle.

## ACKNOWLEDGEMENTS

Matti Leponiemi and Eeva Vakkari kindly helped with field work, and Heini Ali-Kovero and Leena Laaksonen with genetic analyses. Our work was funded by the Academy of Finland (#140990, #135970, #251337, #284666). SH was funded by Finnish Cultural Foundation and Alfred Kordelin Foundation, MI was funded by Betty Väänänen foundation, Societas Pro Fauna et Flora Fennica, Societas Biologica Fennica Vanamo, Suomen Hyönteistieteellinen Seura ry, Helsingin hyönteistieteellinen yhdistys, the Swedish Research Council, and the Bolin Centre for Climate Research, and HH was funded by Kone foundation.

## AUTHOR CONTRIBUTIONS

All authors designed and conceived the study. MI collected the data. SMH and MI analyzed the data, prepared figures and tables, and drafted the manuscript. All authors reviewed drafts and approved the final manuscript.

## DATA ACCESSIBILITY

To be added to final version if the manuscript is accepted. Data will be stored in Dryad.

## APPENDIX

**Table A1.** Number of nests, area and nest density for the studied subpopulations as well as separately for the four parts within the supercolony site. The area of the supercolony site is the area of all the open, and thus potentially habitable, parts of the field. The areas of the individual parts of the supercolony site and the smaller sites are defined as the polygon formed by drawing straight lines between the outermost nests.

| Part | nests | area (ha) | nests/ha |
| --- | --- | --- | --- |
| Supercolony site | 1343 | 5.93 | 226.48 |
| Part I | 281 | 1.11 | 253.78 |
| Part II | 519 | 1.29 | 400.98 |
| Part III | 519 | 1.87 | 277.66 |
| Part IV | 24 | 0.09 | 275.76 |
| Storsand | 16 | 0.63 | 25.26 |
| Lillgård | 7 | 0.02 | 425.81 |
| Storgård | 27 | 1.17 | 23.16 |

**Table A2.** Pairwise comparisons between the factor levels of the fixed effects included in the models.

| **Response** |  |  |  |
| --- | --- | --- | --- |
| Pairwise comparisons | SE | z-value | p |
| **Presense of aggression** |  |  |  |
| **GLMM (binomial)** |  |  |  |
| Neighbor - Distant | 0.401 | 2.231 | 0.026 |
| Neighbor - Outside | 0.432 | 2.887 | 0.004 |
| Neighbor - Allospecific | 0.729 | 6.709 | <0.001 |
| Distant - Outside | 0.406 | 0.866 | 0.386 |
| Distant - Allospecific | 0.708 | 5.639 | <0.001 |
| Outside - Allospecific | 0.718 | 5.075 | <0.001 |
| **Duration of aggression** |  |  |  |
| **GLMM (beta)** |  |  |  |
| Neighbor - Distant | 0.317 | -0.290 | 0.772 |
| Neighbor - Outside | 0.310 | -0.713 | 0.476 |
| Neighbor - Allospecific | 0.299 | 1.111 | 0.267 |
| Distant - Outside | 0.261 | -0.494 | 0.621 |
| Distant - Allospecific | 0.238 | 1.780 | 0.075 |
| Outside - Allospecific | 0.227 | 2.434 | 0.015 |

| Duration of antennation | GLMM (beta) |  |  |
| --- | --- | --- | --- |
| Control - Neighbor | 0.195 | 5.454 | <0.001 |
| Control - Distant | 0.200 | 5.071 | <0.001 |
| Control - Outside | 0.336 | 3.737 | <0.001 |
| Neighbor - Distant | 0.210 | -0.231 | 0.818 |
| Neighbor - Outside | 0.334 | 0.578 | 0.563 |
| Distant - Outside | 0.343 | 0.703 | 0.482 |

Fixed effect levels: Control = introduced ant from the same nest, Neighbor = introduced ant from the same part of the supercolony field, Distant = introduced ant from a different part from the supercolony field, Outside = introduced ant from another field, Allospecific = introduced ant of a different species, *F. exsecta*. The aggression models do not include the level 'Control', for it had no aggression. The model for antennation does not include the level 'Allospecific' due to it having aggression in all but three samples and thus too small a sample size for analysing the non-aggressive behaviors. In all of the models, the nest IDs of both the host and the visitor were included as crossed random effects.

